# Endo-IP and Lyso-IP Toolkit for Endolysosomal Profiling of Human Induced Neurons

**DOI:** 10.1101/2024.09.24.614704

**Authors:** Frances V. Hundley, Miguel A. Gonzalez-Lozano, Lena M. Gottschalk, Aslan N. K. Cook, Jiuchun Zhang, Joao A. Paulo, J. Wade Harper

## Abstract

Plasma membrane protein degradation and recycling is regulated by the endolysosomal system, wherein endosomes bud from the plasma membrane into the cytosol and mature into degradative lysosomes. As such, the endolysosomal system plays a critical role in determining the abundance of proteins on the cell surface, influencing cellular identity and function. Highly polarized cells, like neurons, rely on the endolysosomal system for axonal and dendritic specialization and synaptic compartmentalization. The importance of this system to neuronal function is reflected by the prevalence of risk variants in components of the system in several neurodegenerative diseases, ranging from Parkinson’s to Alzheimer’s disease. Nevertheless, our understanding of endocytic cargo and core endolysosomal machinery in neurons is limited, in part due to technical limitations. Here, we developed a toolkit for capturing EEA1-postive endosomes (Endo-IP) and TMEM192-positive lysosomes (Lyso-IP) in stem cell-derived induced neurons (iNeurons). We demonstrated its utility by revealing the endolysosomal protein landscapes for cortical-like iNeurons and stem cells. This allowed us to globally profile endocytic cargo, identifying hundreds of transmembrane proteins, including neurogenesis and synaptic proteins, as well as endocytic cargo with predicted SNX17 or SNX27 recognition motifs. By contrast, parallel lysosome profiling reveals a simpler protein repertoire, reflecting in part temporally controlled recycling or degradation for many endocytic targets. This system will facilitate mechanistic interrogation of endolysosomal components found as risk factors in neurodegenerative disease.

## Introduction

The endolysosomal system plays a critical role in controlling proteome abundance and quality in eukaryotic cells. First, specific plasma membrane (PM) proteins—referred to here as cargo—undergo the process of endocytosis, where they are concentrated into an inwardly budding membrane structure to form an endosome. Endosomes coordinate three primary protein trafficking systems (1-6): 1) recycling of cargo proteins back to the plasma membrane via sorting into recycling endosomes, 2) retrograde transport of cargo proteins to the Golgi, allowing recycling through the secretory pathway, and 3) trafficking into late endosomal intralumenal vesicles via the ESCRT pathway, ultimately leading to cargo degradation upon maturation into fully degradative lysosomes. These mechanisms collaborate to maintain and dynamically modulate the cell surface proteome while also coordinating intracellular signaling of multiple classes of ligand-activated receptors. Second, the endolysosomal system plays a key role in proteome quality control through the process of autophagy (7). Here, intracellular organelles and proteins are marked for capture within an autophagosome, which then fuse with degradative lysosomes, resulting in degradation of the cargo.

Arguably, among the most demanding settings for spatial control of the endolysosomal system are highly polarized cells such as neurons, which consist of complex dendritic arbors and axons that extend many times the length of the cell body (soma) (8-11). Regulated abundance of PM proteins controls numerous aspects of neuronal function, ranging from axon guidance to the identity and function of pre- and post-synaptic compartments. The distinct dendritic and axonal cohorts of PM proteins are controlled by the axon initial segment and regulated endocytic degradation (12-14). Moreover, organelle quality control via autophagy especially within axons relies on the formation of endolysosomes which mature to form lysosomes that are capable of fusion with autophagosomes distally (15). Such autolysosomes are subsequently trafficked back to the soma and become sufficiently acidic for support of cargo degradation during transit to the soma (16). The importance of endolysosomal trafficking for neuronal health is indicated by the number and diversity of genes in the pathway that are risk variants across neurodegenerative diseases, ranging from Parkinson’s to Alzheimer’s diseases, including LRRK2, VPS35, BIN1, and PSEN1/2 (17-21). Defects in such pathways may alter the location and abundance of critical PM proteins, as demonstrated for the VPS35 pathway (22).

A current limitation in the dissection of endolysosomal function is the lack of approaches for analysis of specific organelles within the endolysosomal system in neuronal systems. Here, we report the development of a toolkit for profiling a population of early/sorting endosomes, as well as lysosomes (23), in human ES cell (hESC)-derived cortical-like iNeurons. We target a sub-population of early endosomes marked by endogenously tagged EEA1 (<u>E</u>arly <u>E</u>ndosome <u>A</u>ntigen 1) protein, which represents one of the earliest populations formed after endocytic vesicle uncoating. Given the proximity of this population to uncoating, this “Endo-IP” approach (24) allows detection of the largest repertoire of endocytic cargo, prior to degradation during maturation to lysosomes. Our work revealed broad remodeling of endocytic cargo during *in vitro* neurogenesis, reflecting programmed expression of a diverse repertoire of PM protein involved in numerous neuronal functions. In contrast, lysosomes display a much simpler proteome rich in hydrolytic enzymes, reflecting both cargo sorting for retrograde transport or degradation during endolysosomal maturation. Finally, we present a functional landscape of endocytic cargo, as well as a compendium of candidate cargo sorting motifs and structural predictions for two major sorting motifs linked with SNX27-Retromer and SNX17-Retriever (21, 25-28). This work provides the tools for mechanistic interrogation of pathways controlling cargo selection and the impact of neurodegenerative disease risk alleles on this process.

## Results and Discussion

### Genetic tagging of EEA1 and TMEM192 in human stem cells

In order to purify endolysosomal compartments from human neurons, we adopted the Endo-IP approach using EEA1 protein as an affinity handle, incorporating three copies of the Flag epitope (24) (**Fig. 1A**). Unlike gradient fractionation of endosomes which captures many classes of endosomes (29), the EEA1 affinity handle allows capture of a small population of endosomes that is expected to be enriched in newly endocytosed cargo. To avoid any effects of Flag-EEA1 overexpression, we endogenously tagged both EEA1 copies using CRISPR-Cas9 in hESCs harboring an inducible AAVS1-NGN2 neurogenesis driver (30-32) (**Fig. S1A**). In parallel, we similarly edited Flag-EEA1 in hESCs previously engineered to contain the Lyso-IP affinity handle—a triple HA epitope fused with the C-terminus of TMEM192 (in heterozygous form, see **Supplemental Methods**) (**Fig. 1A and Fig. S1B**). These cell lines were validated by sequencing and immunoblotting of cell extracts from hESCs and day 21 iNeurons (**Fig. S1A-D**). We refer to singly and doubly tagged cells as “H9-E” and “H9-EL”, with “E” referring to Endo-IP and “EL” referring to Endo- and Lyso-IP.

**Fig. 1:**
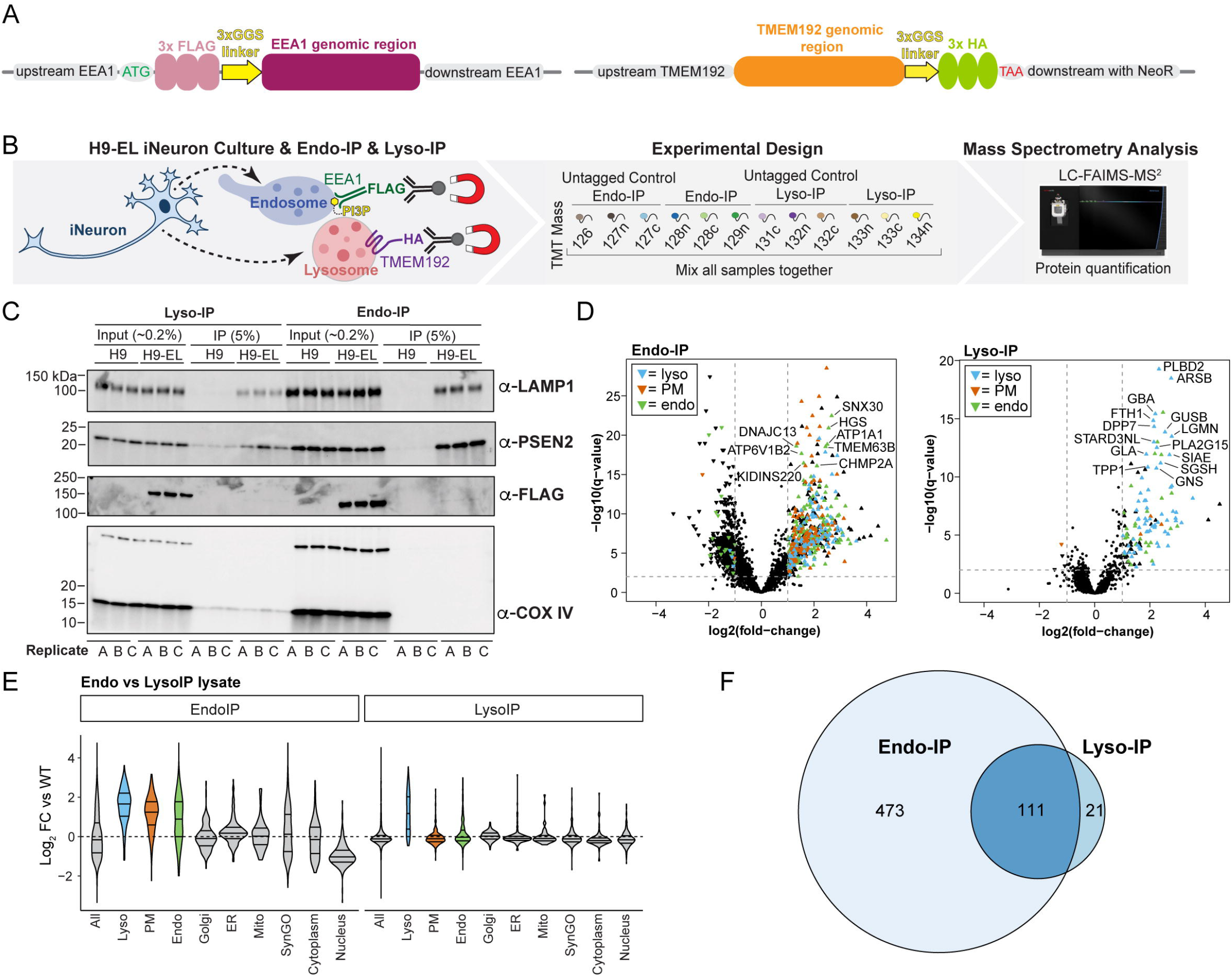
Toolkit and validation of Endo-IP and Lyso-IP for organelle proteomics in hESC-derived iNeurons. **(A)** Diagram of endogenous tagging of the N-terminus of EEA1 and the C-terminus of TMEM192 with 3XFlag and 3XHA tags using CRISPR-Cas9. **(B)** Schematic of TMT-based proteomic analysis of Endo-IP and Lyso-IP from H9-EL-derived day 21 iNeurons. EEA1-positive endosomes are captured with *α*-FLAG conjugated to magnetic beads and TMEM192-positive lysosomes are similarly captured with *α*-HA antibodies. Purified EEA1-positive or TMEM192-positive membranes were eluted from the beads with a mild detergent, and proteins were digested with LysC and trypsin. The resulting peptides were labeled with 12-plex TMT and analyzed by mass spectrometry. **(C)** Immunoblotting of the indicated extracts or immunoprecipitation (IP) samples using *α*-LAMP1 and *α*-PSEN2 to indicate endolysosomes and *α*-Flag to detect Flag-EEA1. *α*-COXIV was used as a control for non-specific capture of mitochondria. “A, B, and C” refer to replicate experiments. The percentage of input and IP loaded on the gel are indicated. **(D)** Volcano plots of log_2_ fold change (FC) for tagged versus untagged (control) iNeurons versus -log10(q-values) are plotted. Proteins annotated as lysosomal (cyan), plasma membrane (orange) or endosomal (green) are indicated. **(E)** Violin plots of log_2_ FC relative to WT (untagged control) iNeurons for various groups of proteins. **(F)** Venn diagram for overlap of proteins enriched in the Endo-IP and Lyso-IP from iNeurons.

To ensure proper localization of Endo- and Lyso-IP tags, we performed immunofluorescence in combination with confocal microscopy on day 21 iNeurons. As expected, α-Flag and α-EEA1 signals displayed a very high degree of colocalization (Mander’s coefficient >0.85) (**Fig. S1E**,**F**). Consistent with EEA1 colocalization with a subset of early/sorting endosomes, small mean Mander’s coefficients of 0.15-0.28 were observed for α-RAB5 and α-FLAG or α-RAB5 and α-EEA1 in both H9-E and H9-EL cell lines (**Fig. S1E**,**F**). The extent of colocalization was slightly, but not significantly reduced in H9-E and H9-EL tagged cells compared to WT untagged controls (mean Mander’s coefficients of 0.2-0.3 vs 0.4) **(Fig. S1E**,**F**). For Lyso-IP tagged iNeurons, TMEM192-HA puncta partially co-localized with the lysosomal marker LAMP1 with Mander’s coefficients in the range of both α-HA with α-TMEM192 and α-LAMP1 with α-TMEM192 (0.3-0.45). Mander’s coefficients were not significantly different between Lyso-IP-tagged and untagged iNeurons (mean Mander’s coefficients of 0.3-0.35 for both) **(Fig. S1E**,**F**). The extent of RAB5 and EEA1 colocalization was in agreement with the broad range reported previously in HEK293 (24) and HeLa (33) cells. Cell line variability in the distribution of EEA1 between early/sorting endosomes and late endosomes has been observed previously (33). Taken together, these data indicate the validity of these cells for analysis of EEA1-positive populations of endosomes and of TMEM192-positive lysosomes.

### Benchmarking Dual-tagged EL cells for Organelle Proteomics

To initially benchmark H9-EL cells for organelle IP, we converted control (untagged H9) and H9-EL cells to day 21 iNeurons in biological triplicate and performed Endo- and Lyso-IPs followed by either immunoblotting or 12-plex Tandem Mass Tagging (TMT)-based mass spectrometry (MS) (**Fig. 1B**). Immunoblotting of Endo- or Lyso-IP samples demonstrated selective enrichment of LAMP1 and PSEN2, with no enrichment of COX IV as a mitochondrial marker (**Fig. 1C**). Proteomic analysis revealed that Lyso-IP samples were substantially enriched in known lysosomal proteins, including catabolic enzymes (GNS, GUSB, GBA, GLA), among many others (**Fig. 1D and E and Fig. S2A, Dataset S1**). Indeed, of the 132 proteins with Log_2_ FC>1.0, 104 were known lysosomal proteins and 7 others were annotated as secreted or plasma membrane-localized (**Fig. 1D and Fig. S2D**). Moreover, analysis of several subcellular compartments demonstrated a high degree of selectivity in the enrichment of lysosomal proteins (**Fig. 1E and Fig. S2B**). Of the 91 proteins enriched in Lyso-IPs from HEK293 cells, 56 were enriched in iNeurons (**Fig. S2C**). In contrast with Lyso-IP samples, the overall number of proteins enriched in the Endo-IP of iNeurons was ∼4 times larger than with Lyso-IP and was substantially populated with PM proteins in addition to numerous proteins annotated as endosomal or lysosomal (**Fig. 1D-F and Fig. S2A**,**B, Dataset S1**). As expected, proteins found in common in both Endo-IP and Lyso-IP are typically core components of the endolysosomal system, including BORCS6, LAMTOR2, and ATP13A2 (**Fig. 1F and Fig. S2D**). These results indicate the feasibility of endosome and lysosome proteome profiling in iNeurons and the capacity of Endo-IP to capture protein cargo.

### Proteomic Landscape of Endosomal Cargo During *in vitro* Neurogenesis

Our previous studies have demonstrated large-scale proteome remodeling during NGN2-driven conversion of hESCs to iNeurons *in vitro*, including the induction of numerous factors that drive the neurogenesis program (31). A total of 476 endolysosome-related proteins (see **Supplemental Methods**) were categorized in clusters based on their abundance profile during a 12-day differentiation, revealing that most proteins dramatically increased in abundance during neurodifferentiation (**Fig. 2A**). As expected, many proteins annotated in Synaptic Gene Ontology (SynGO)—an evidence-based and curated database for synaptic proteins (34)—also displayed similar alterations in abundance during differentiation (**SI Appendix Fig. S3A**). This includes markers for synaptic vesicles (SYP), postsynaptic receptor (GRIA2), dendrites (MAP2) and adhesion (NCAM1) (**Fig. 2B**).

**Fig. 2:**
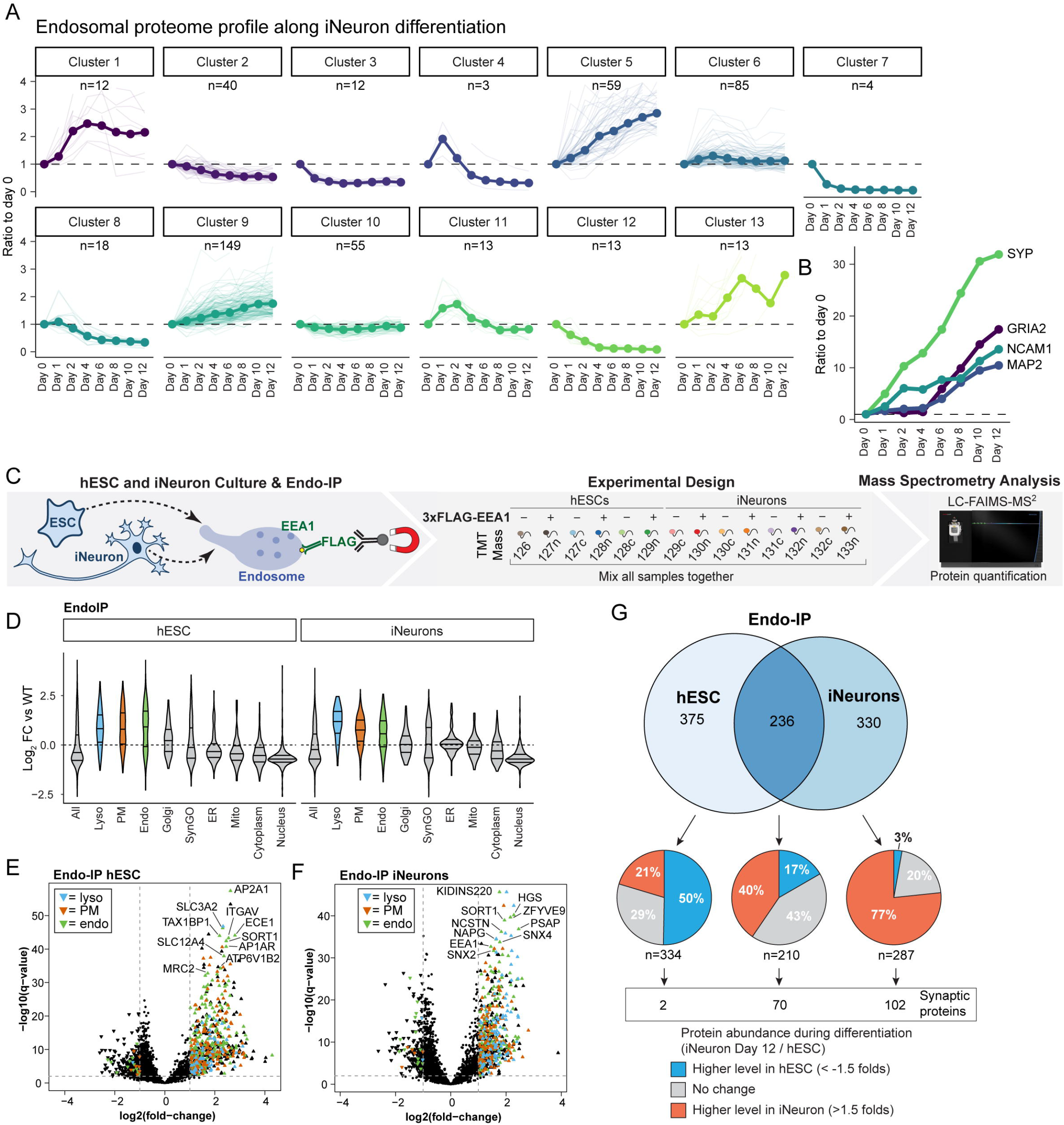
Proteomic Landscape of EEA1-positive Endosomes in hESCs and iNeurons. **(A)** Alterations in endosomal proteins during *in vitro* differentiation of hESCs to iNeurons, displayed as clusters identified based on patterns of expression, based on data from our previously reported TMT-based proteomic analysis of H9 cells using the NGN2 driver (31). **(B)** Analogous to panel A but for 4 markers of neurogenesis (SYP, GRIA2, NCAM1, and MAP2). **(C)** Schematic of our approach for quantitative analysis of the hESC and day 21 iNeuron endosomal proteome using the Endo-IP method for H9-E cells, and parental H9 cells as a control in triplicate or quadruplicate. Purified EEA1-positive membranes were eluted from the beads with a mild detergent, and proteins were digested with LysC and trypsin. The resulting peptides were labeled with 14-plex TMT and analyzed by mass spectrometry. **(D)** Violin plots of log_2_ FC relative to WT (untagged control) cells for various groups of proteins found in Endo-IPs from hESCs or day 21 iNeurons. **(E and F)** Volcano plots of log_2_ FC for tagged versus untagged (control) hESCs (panel **E**) or iNeurons (panel **F**) versus -log10(q-values) are plotted. Proteins annotated as lysosomal (cyan), plasma membrane (orange) or endosomal (green) are indicated. The dashed lines indicate the log_2_ FC cut-off of 1.0 and -log10(q-value) cut-off of 2 (q-value of 0.01). **(G)** Comparison of proteins identified by Endo-IP in hESCs and day 21 iNeurons (upper Venn diagram). Pie charts display the proportion of identified proteins with abundance changes in either the hESC or iNeuron state, based on (31). The number of proteins present in SynGO are indicated.

To identify endocytic factors and neuron-specific cargo, we performed TMT-MS on triplicate or quadruplicate Endo-IPs from H9 control or H9-E cells in either the stem cell state or iNeurons at day 21 of differentiation (**Fig. 2C**). As expected, immunoblotting of iNeuron Endo-IP samples revealed enrichment for LAMP1 as an endolysosomal marker, but not Golgi (GOLGA1) or ER (CALR) markers (**Fig. S3B**). The reproducibility of the method was assessed by comparing two independent Endo-IP datasets, which revealed extensive overlap of enriched proteins identified and reproducible protein levels (**Fig. S3C**,**D**). Consistent with the results described above, Endo-IPs from both hESC and iNeuron populations displayed enrichment of numerous endolysosomal and plasma membrane proteins, but no enrichment in other cellular compartments (Log_2_FC ≥ 1.0, q-value ≤ 0.01) (**Fig. 2D-F and Fig. S3E**). From the 611 and 566 proteins enriched in Endo-IPs in hESCs and iNeurons, respectively, 236 were common to both cell types (**Fig. 2G**). These included nutrient transporters (ECE1, SLC3A2), subunits of sorting complexes (AP2A1, VPS35, SORT1, SNX4, SNX2, and NAPG), v-ATPase subunits (ATP6V1B1, ATP6V1E1), signal transduction factors (ZFVYE9), autophagy pathways components (TAX1BP1, GABARAP, GABARAPL1, and GABARAPL2), and other established endosomal factors or cargoes (PSAP, EEA1, HGS, and NCSTN) (**Fig. 2G and Dataset S2**). Enriched proteins that were selective for one or the other cell state tended to correlate with the relative levels of expression; 79% of hESC-selective proteins had higher abundance (50%) or were of equal abundance (29%) in hESCs while 97% of iNeuron-selective proteins had higher abundance (77%) or were of equal abundance (20%) in iNeurons (**Fig. 2G**). Similarly, 102 synaptic proteins were identified exclusively in iNeurons, while only 2 and 70 were identified only in hESCs or both hESCs and iNeurons, respectively. Taken together, these data indicate that hESCs and iNeurons have non-overlapping EEA1-positive endosomal proteomes.

### Repertoire of Receptor and Synaptic Protein Cargo in iNeurons

To characterize the endolysosomal proteins identified in iNeurons, we first organized Endo-IP enriched proteins into broad functional categories, encompassing many core endolysosomal functions, including sorting (Retromer, ESCRT), membrane fusion (SNAREs), positioning (BORC), and numerous core endolysosomal proteins (**Fig. 3A**). Many of these proteins, indicated by asterisks, are also present in SynGO, consistent with known roles for endosomal vesicle fusion and vesicle positioning within the synaptic compartment (**Fig. 3B**). SynGO analysis also revealed a significant enrichment of additional synaptic proteins in the iNeuron Endo-IP not previously linked with core endosomal functions, including several categories of pre- and post-synaptic sub-cellular locations, and functional categories (e.g. synaptic organization, signaling and transport) (**Fig. 3B and C and Fig. S4A**,**B**). Further mining of the synaptic proteins revealed numerous proteins linked with synaptic endosomes and synaptic membranes that most likely represent endocytic cargo, featuring diverse transmembrane receptors and adhesion proteins (**Fig. 3C**).

**Fig. 3:**
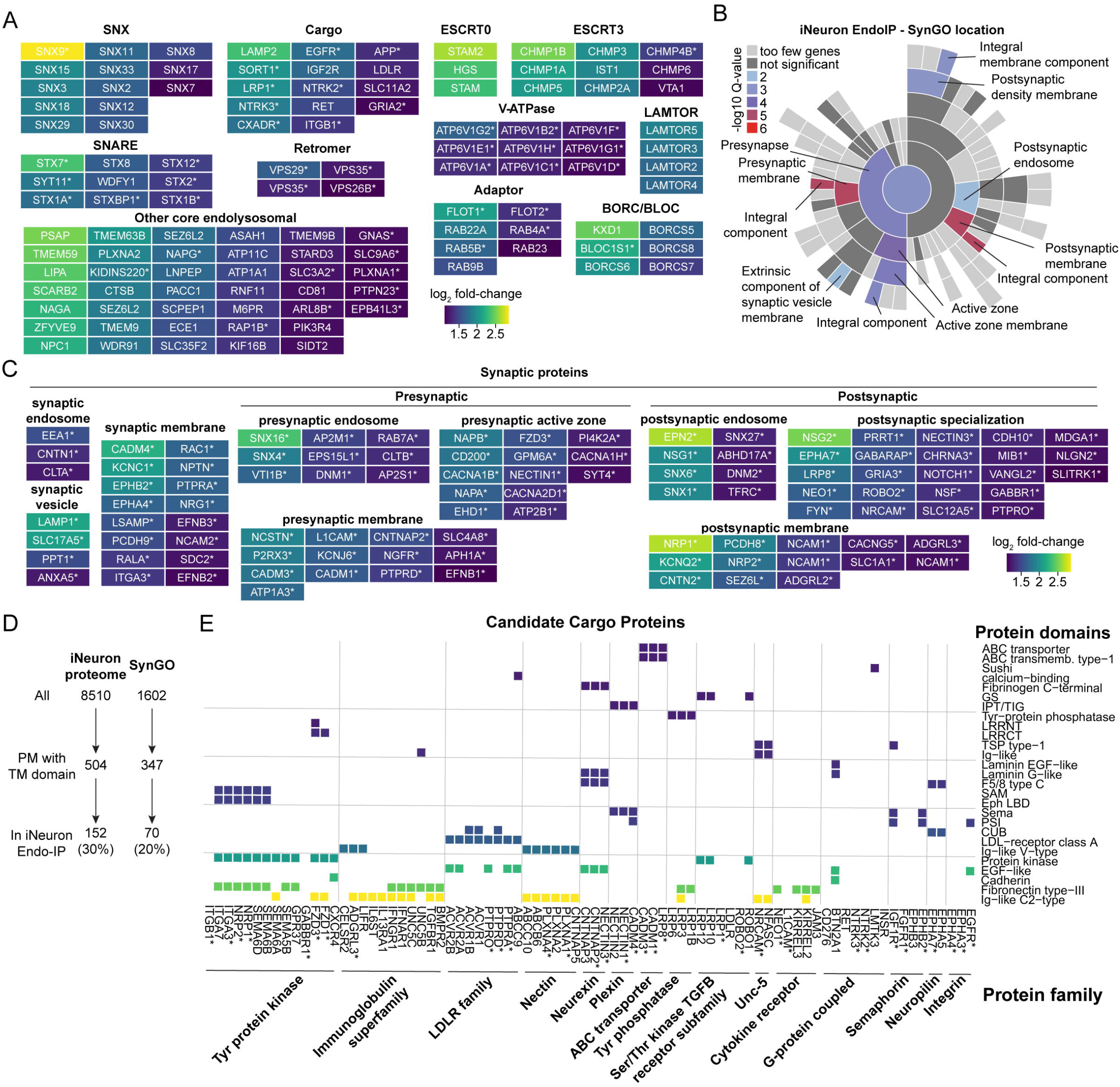
Repertoire of Receptor and Synaptic Protein Cargo in iNeurons. **(A)** Functional categories of endosomal proteins identified by Endo-IP in day 21 iNeurons. Color code refers to the extent of enrichment in Endo-IP, as indicated by the scale bar. Proteins in these categories that are also present in SynGO are indicated with an asterisk. **(B)** Enrichment of iNeuron Endo-IP proteins based on SynGO location. Color code corresponds to the log_10_ of q-value for individual categories of SynGO proteins. **(C)** Categories of synaptic proteins identified as being enriched in Endo-IPs from day 21 iNeurons. **(D)** Relative enrichment of iNeuron proteome (31) and SynGO proteome within the Endo-IP from day 21 iNeurons. **(E)** Classification of a sub-set of transmembrane proteins and their associated extracellular domains and protein families identified as enriched in Endo-IPs from day 21 iNeurons. Colored squares reflect the presence of the indicated domain within individual proteins. Proteins are organized based on their presence within the indicated protein family. An asterisk indicates that the protein is present in SynGO.

To further understand the repertoire of candidate cargo in iNeurons, we examined all proteins harboring one or more transmembrane segments which are also annotated as localized in the cell membrane based on Uniprot. We found 504 and 347 of such proteins in iNeurons and SynGO, of which 152 and 70 were also identified in Endo-IP, respectively (**Fig. 3D**). This indicates that 30% of all plasma membrane proteins in iNeurons and 20% of plasma membrane proteins with annotated synaptic localization are candidate cargo for endocytosis. We next identified 241 candidate cargo specifically in our Endo-IP in iNeurons, applying the same transmembrane and cell membrane localization criteria (**Dataset S2**), and further categorized candidate cargo by their annotated domains and protein families (**Fig. 3E and Fig. S4C**). Of the 80 candidate cargo containing both an annotated domain and protein family (including 34 in SynGO), more than a dozen types of extracellular domains were represented. These included Ig-like C2-type, fibronectin, EGF-like and protein kinase domains, found in members of the Nectin, low density lipid receptor (LDLR), Neurexin, immunoglobulin, and tyrosine protein kinase families of proteins (**Fig. 3E**). Additionally, 89 further candidate cargo (including 24 in SynGo) with annotations for either a domain or a protein family, including channels, transporters, peptidases and tetraspanin proteins, were identified (**Fig. S4C**). Given the kinetics of protein recycling or degradation upon endocytosis (on the order of minutes), the diversity of proteins identified with Endo-IP enrichment suggests extensive ongoing remodeling of the plasma membrane proteome during in vitro neurogenesis.

### Matching Candidate Cargo with Sorting Motifs

Early endosomes function as sites of cargo sorting for recycling to the plasma membrane or Golgi, and also further mature to late endosomes, whereas cargo destined for turnover are internalized into intralumenal vesicles for subsequent degradation in the lysosome (4). Retromer and Retriever function as sorting complexes in conjunction with a variety of sorting nexins, including SNX27 and SNX17, respectively (22, 26, 27, 35). Mutations in the VPS35 subunit of Retromer have been linked with Parkinson’s disease, while mutations in the CCDC22 subunit of Commander (Retriever plus the CCC complex) are found in Ritscher-Schinzel syndrome (21, 36). To initially examine the relationship between candidate cargo identified in the iNeuron system and known SNX17 and/or SNX27 cargo, we used previously reported data sets (22, 26) to ask to what extent our candidate cargo were among proteins whose abundance on the plasma membrane was altered in HeLa cells depleted of SNX17 or SNX27. For this purpose, we employed the 241 proteins in our iNeuron dataset that contained at least one transmembrane domain and that were annotated in Uniprot as being localized in the cell membrane (**Dataset S3**). The majority (75%) of iNeuron proteins were not previously identified in the SNX17/SNX27 studies, potentially reflecting cell type diversity, while 16 to 21 proteins (6.5-8.5%) were common to two or three datasets (**Fig. 4A**). The finding that many candidate cargo in iNeurons were absent from known cargo led us to examine these proteins for the presence of sorting motifs.

**Fig. 4:**
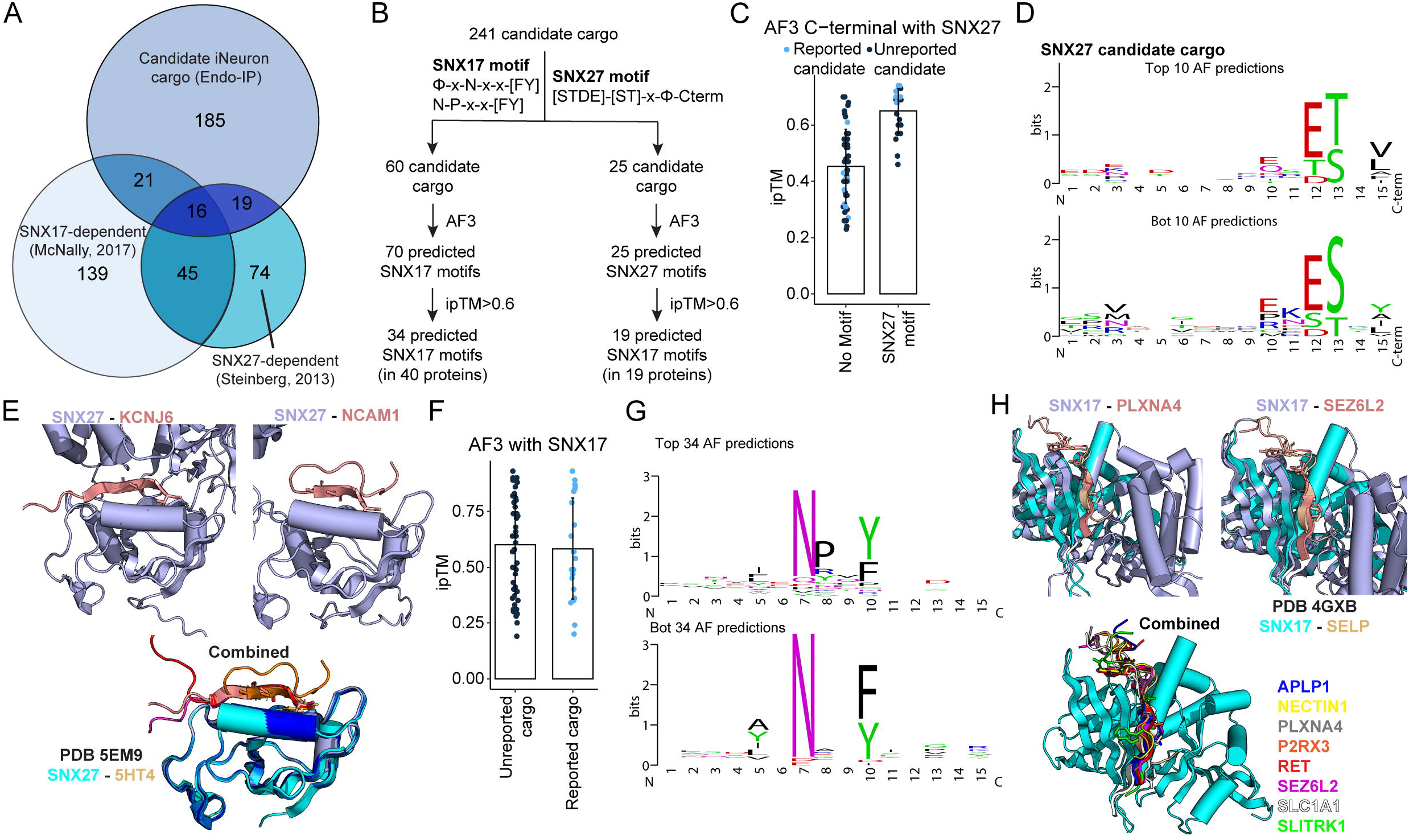
Matching candidate cargo with SNX17 and SNX27 sorting motifs. **(A)** Comparison of the proteins identified by Endo-IP in day 21 iNeurons with proteins whose cell surface levels depend on SNX17 or SNX27 or proteins that associate with SNX27 (22, 26).The number of proteins in each category is indicated. **(B)** Scheme summarizing the search for SNX17 or SNX27 sorting motifs in 241 candidate cargo. Candidate motifs were subjected to predictions using AF3 and the number of proteins with ipTM scores >0.6 indicated. **(C)** Plot of ipTM values for AF3 predictions involving proteins with or without candidate SNX27 motifs. Light blue dots represent proteins reported as candidates for SNX27 cargo. **(D)** Motif plots for Top 10 and Bottom (Bot) 10 AF3 predictions. **(E)** Structural predictions for candidate iNeuron cargo and the PDZ domain of SNX27. C-terminal motifs are shown in salmon and SNX27 in light blue. **(F)** Plot of ipTM values for AF3 predictions involving candidate proteins with SNX17 motifs. Light blue dots represent proteins reported as candidate SNX17 cargo. **(G)** Motif plots for Top 34 and Bottom (Bot) 34 AF3 predictions. **(H)** Structural predictions for candidate iNeuron cargo and the FERM domain of SNX17. SNX17-motif peptide predictions are in light blue (SNX17) and salmon (motifs). The structure of SNX17-SELP (27) (PDB: 4GXB) is overlayed in cyan and brown.

SNX27 uses its PDZ domain to bind [S/T]-x-Φ-C-terminal motifs in cargo (in which Φ represents any hydrophobic residue as the C-terminal residue in the candidate protein) (22, 25, 37, 38). This core interaction is frequently preceded by one or more acidic residues or phosphorylated Ser/Thr residues in a manner that enhances binding (28). In contrast, SNX17 uses its FERM (4.1/ezrin/radixin/moesin) domain to bind to ΦxNxx[YF] and related motifs (where Φ is a hydrophobic amino acid) in cargo (27). Motif searches revealed 25 and 54 candidate cargo containing potential SNX27 and/or SNX17 interaction motifs, respectively (**Fig. 4B**). For the 25 SNX27 candidates, we screened their C-termini as 15 amino acid peptides for interaction with the PDZ domain of SNX27 using AF3 (39). In parallel, we screened a set of 47 potential SNX17 cargo that lacked an apparent SNX27 binding motif as controls for AF3 predictions (**Fig. 4B**). We identified 19 candidate cargo with AF3 ipTM scores >0.6 (**Fig. 4C and Dataset S3**). In contrast, the majority of control sequences lacking an SNX27 motif had ipTM scores <0.6. Importantly, of the candidate SNX27 cargo that were also identified previously in HeLa cells, the majority had ipTM scores of >0.6 (**Fig. 4C**). As expected, the motifs identified among the top 10 and bottom 10 AF3 predictions conform to the canonical SNX27 sorting motif (**Fig. 4D**). Examples of SNX27 PDZ domain-candidate sorting motif predictions for KCNJ6, KCNJ12, NCAM1, and NCAM2 are shown independently and together with the previously determined structure of SNX27 and 5HT4 (PDB 5EM9) (28) (**Fig. 4E and Fig. S5A**).

Similarly for candidate SNX17 cargo, a total of 70 potential cargo sorting motifs in 60 candidate proteins were screened by AF3, leading to the identification of 34 candidate SNX17 sorting motifs in 40 proteins with ipTM scores >0.6 (**Fig. 4B and F and Dataset S3**). The distribution of ipTM scores for known and previously unidentified candidate cargo were similar, with many known cargo having ipTM scores less than 0.6 (**Fig. 4F**). As expected, the motif identified for the top scoring cargo fit the consensus for SNX17 FERM binding proteins (**Fig. 4G**). The predicted structures for several candidate cargo closely matched the previously determined structure for SNX17 and SELP (PDB: 4GXB) (27) (**Fig. 4H and Fig. S5B**). Taken together, these data uncovered novel neuronal-specific cargo in human neurons and showcase how Endo-IP can be used to enrich and identify candidate cargo for specific sorting pathways.

## Conclusions

Stem cell derived neurons have emerged as important tools for understanding the basic cell biology and functions of genes linked with neurogenesis. Multiple lineage drivers have been developed, affording the potential to create distinct classes of iNeurons with unique properties (32, 40-42). iNeurons maintain many features of animal-derived neurons, including intact axonal and dendritic trafficking pathways, and reliance on organelle quality control systems such as autophagy to remove damaged or superfluous organelles (16, 43, 44). In addition, efforts to generate diverse mutations linked with neurodegenerative diseases have been initiated, with the long-term goal of understanding how risk alleles linked with specific diseases alter critical trafficking and quality control pathways (45). With these advances in mind, we set out to develop tools to isolate early endosomes and lysosomes using genetically tractable hESC-derived cortical-type iNeurons. We demonstrate the utility of endogenously-tagged EEA1 and TMEM192 genes with Flag and HA epitopes, respectively, for isolation of early endosomal and lysosomal populations from day 21 cortical-type iNeurons. Thus, overexpression of tagged affinity handles, as employed previously (23, 24), is not required for effective capture of organelles.

Our studies reveal a diverse array of candidate endocytic cargo identified by Endo-IP, including many classes of single and multi-pass transmembrane proteins functioning in a variety of pathways associated with neurogenesis. Through systematic motif analysis coupled with AF3-based predictions, we provide a compendium of candidate cargo for SNX27- and SNX17-based sorting machinery, which function with Retromer and Retriever complexes, respectively (4, 21). Interestingly, our approach identified several candidate cargo for SNX17 or SNX27, including several proteins linked with neurogenesis and or neuronal function (e.g. PLXNA4, P2RX3, RET, SEZ6L2, SLC1A1, SLITRK1) (**Fig. 4H and Fig. S5A**,**B**). Further directed studies in the context of sorting mutants are required to validate specific candidate motifs and associated machinery, and any spatial aspects of endocytic selectivity (12), but our analysis provides a broad landscape for such studies. Moreover, we envision that distinct sets of cargo will be identified in distinct types of iNeurons, in the context of activation, or in the presence of glial cells which promote the formation of functional synaptic compartments. Additionally, our compendium of candidate cargo will facilitate studies designed to understand spatial control of endocytosis. Taken together, our toolkit enables profiling diverse aspects of the endolysosomal components to further understand the system in physiological and neurodegenerative conditions.

## Materials and Methods

Detailed procedures related to cell culture, gene editing, biochemical procedures, mass spectrometry, microscopy and flow cytometry are described in *SI Appendix*, Materials and Methods. Statistical analyses were performed using MSstats and ggplot. All error bars represent standard error of the mean (SEM), and statistical significance was determined by T-tests, as specified in the corresponding figure legends.

## Data and Material Availability Statement

Uncropped blots, light microscopy images, raw proteomic data and AF3 prediction PDB files will be deposited at Zenodo.org upon publication.

## Acknowledgments

We thank members of the Harper lab for feedback. This work was funded by Aligning Science Across Parkinson’s (ASAP) (J.W.H.), NIH R01NS110395 and R01NS083524 (J.W.H.), the Warren Alpert Foundation (J.W.H.), NIH R01 GM132129 (J.A.P.), a Robin Reed Memorial Postdoctoral Fellowship (F.V.H.), and a Rubicon Postdoctoral Fellowship (M.A.G.-L.). Michael J. Fox Foundation administers the grant ASAP-000282 on behalf of ASAP and itself. For the purpose of open access, the author has applied for a CC-BY public copyright license to the Author Accepted Manuscript (AAM) version arising from this submission. We acknowledge the Core for Imaging Technology and Education (CITE, Harvard Medical School) for imaging assistance.

