## Supplemental Methods and Supplemental Figures for "Endo-IP and Lyso-IP Toolkit for Endolysosomal Profiling of Human Induced Neurons"

**This PDF file includes:**

Materials and Methods
Figures S1 to S5
Legends for Datasets S1 to S3
Supplemental References

**Other supporting materials for this manuscript include the following:**

Datasets S1 to S3

### Supporting Information Text

### MATERIALS and METHODS

#### Reagents

The following chemicals and reagents were used: Nunc Nunclon Delta cell culture dishes (Thermo Scientific, 140675, 167008, 168381, 172931); Corning Matrigel Matrix (Corning, 354230); DMEM/F12 (Gibco, 11330057); Neurobasal Medium (Thermo Scientific, 21103049); non-essential amino acids (NEAAs, Gibco, 11140050); GlutaMAX (Gibco, 35050061); N-2 supplement (Gibco, 17502048); neurotrophin-3 (NT3; PeproTech, 450-03); brain-derived neurotrophic factor (BDNF; PeproTech, 450-02); B27 (Gibco, 17504001); Y27632 dihydrochloride (ROCK inhibitor; PeproTech, 1293823); Accutase (StemCell Technologies, 07922); FGF2-G3 (in-house); human insulin (Santa Cruz Biotechnologies, sc-360248); transforming growth factor-β (TGF-β; PeproTech, 100-21); holo-transferrin human (Sigma-Aldrich, T0665); sodium bicarbonate (Sigma-Aldrich, S5761-500G); sodium selenite (Sigma-Aldrich, S5261-10G); doxycycline (Clontech Labs, 631311); UltraPure 0.5 M EDTA (Invitrogen, 15575020); 16% paraformaldehyde (Electron Microscopy Science, 15710); CloneR (StemCell Technologies, 05889); 24-well glass bottom plates (Cellvis, P24-1.5H-N); NEB Next Ultra II Q5 Master Mix (New England Biolabs, M0544L); Cas9-NLS (QB3 MacroLab, UC Berkeley), MiSeq Reagent Nano Kit v2 (300 cycles; Illumina, MS-103-1001); GeneArt Precision gRNA synthesis kit (Thermo Scientific, A29377); Pierce Anti-HA Magnetic Beads (Thermo Scientific, 88837); Anti-FLAG M2 Magnetic Beads (Sigma-Aldrich, M8823); IGEPAL CA-630 (Sigma-Aldrich, I8896); S-Trap micro columns (Protifi, C02-micro-80); Triethylammonium bicarbonate buffer (TEAB, Sigma-Aldrich, T7408); sodium dodecyl sulfate (SDS; Bio-Rad, 1610302); Precision Plus Protein Kaleidoscope pre-stained protein standard (BioRad, 1610375); TMTpro 16plex Set (Thermo Scientific, A44520); protease inhibitor cocktail (Roche, 4906845001); tris(2-carboxyethyl)phosphine (TCEP; Gold Biotechnology, 51805-45-9); 2-chloroacetamide (Sigma-Aldrich, C0267); trypsin (Promega, V511C); Lys-C (Wako Chemicals, 129-02541); 50% hydroxylamine solution (Sigma-Aldrich, 438227), High-pH Reversed-Phase Peptide Fractionation Kit (Thermo Scientific, 84868), Bio-Rad Protein Assay Dye (Bio-Rad, 5000006); Empore SPE Disks C18 (Sigma-Aldrich, 66883-U);

The following primary antibodies were used (1:500-1:1,000 for immunoblotting or 1:100-1:200 for immunofluorescence): FLAG (Sigma-Aldrich, F1804), FLAG (Thermo Scientific, MA1-91878), HA (Cell Signaling Technology, 2367), HA (Roche, 11867423001), actin (Santa Cruz Biotechnology, SC-69879), HSP90 (Santa Cruz Biotechnology, SC-69703), RAB5 (Cell Signaling Technology, 3547), LAMP1 (Cell Signaling Technology, 9091), LAMP1 (Cell Signaling Technology, 15665), EEA1 (Cell Signaling Technology, 3288), EEA1 (Thermo Scientific, GT10811), TMEM192 (Proteintech, 28263-1-AP), PSEN2 (Abcam, ab51249), CALR (Proteintech, 10292-1-AP), GOLGA1 (Proteintech, 12640-1-AP), COXIV (Cell Signaling Technology, 4850), VPS35 (Santa Cruz Biotechnology, SC-374372). The following secondary antibodies were used (1:5,000-1:10,000 for immunoblotting, 1:400 for immunofluorescence): goat anti-rabbit immunoglobulin-G (IgG)-horse radish peroxidase (HRP) conjugate (BioRad, 1706515); goat anti-mouse IgG HRP conjugate (BioRad, 1706516); goat anti-mouse IgG (H+L) highly cross-adsorbed, Alexa Fluor 488 conjugate (Thermo Scientific, A-11029); goat anti-rabbit IgG (H+L) cross-adsorbed, Alexa Fluor 488 conjugate (Thermo Scientific, A-11034); goat anti-rabbit IgG (H+L) cross-adsorbed, Alexa Fluor 647 conjugate (Thermo Scientific, A-21244); and goat anti-rat IgG (H+L) cross-adsorbed, Alexa Fluor 647 conjugate (Thermo Scientific, A-21247),

#### Cell line construction and maintenance

Gene edited human embryonic stem cells (hESCs, H9, WiCell Institute) were cultured as previously described (https://dx.doi.org/10.17504/protocols.io.j8nlkoq56v5r/v1) (1). Cells were maintained in E8 medium on plates coated with Matrigel (coated at approximately 8.3 μg/cm$^2$) and split with 0.5 mM EDTA in DPBS.

hESCs with the AAVS1-TRE3G-NGN2 driver were differentiated into iNeurons as previously described (https://dx.doi.org/10.17504/protocols.io.x54v9p8b4g3e/v1) (2, 3). Briefly, on differentiation day 0, hESCs were plated in ND1 medium (DMEM/F12, N-2, human 10ng/mL BDNF, 10 ng/mL NT3, NEAA, 0.2μg/mL human laminin) supplemented with 2μg/mL doxycycline and 10 μM Y27632 (ROCK inhibitor). The next day the medium was exchanged to ND1 supplemented with 2μg/mL doxycycline but without Y27632. The following day, the medium was replaced with

ND2 (neurobasal medium, B27, GlutaMAX, 10ng/mL BDNF, 10ng/mL NT3) supplemented with 2μg/mL doxycycline. Cells were replated on day 4, 5, or 6, depending on density, and for immunofluorescence experiments only (not Endo-IP or Lyso-IP experiments), cells were replated again on day 12-14 into Matrigel-coated 24W glass-bottomed dishes. From day 10, doxycycline was removed from the ND2. From the replating at day 4, 5, or 6 until the experimental day (day 21-23), 50% of the medium was replaced with fresh ND2 every other day.

**CRISPR–Cas9 gene editing**

hESCs (H9 AAVS1-TRE3G-NGN2 RRID:CVCL_C4EK or H9 AAVS1-TRE3G-NGN2 TMEM192-HA, see dx.doi.org/10.17504/protocols.io.e6nvwdjd7lmk/v1) were endogenously tagged using CRISPR-Cas9 (4). Cells were electroporated with a mixture of 0.6µg guide RNA (sgRNA) and 3µg Cas9-NLS (QB3 MacroLab, UC Berkeley) using a Neon transfection system (Thermo Scientific) as previously described (https://dx.doi.org/10.17504/protocols.io.ewov1qykkgr2/v1). The N-terminus of EEA1 was endogenously tagged with 3xFLAG using the following ultramer repair template: gcagggtctggagagtcaccgcggcggcgccgggtggtggttaaaccatgGACTACAAAGACCATGACGGTGATTA TAAAGATCATGACATCGATTACAAGGATGACGATGACAAGGGCGGATCCGGGGGAAGCGGC GGATCCttaaggaggattttacagagggtaagagagtgaaaccgtctttctggcagcacgtg (homology arms in lower case). The EEA1 N-terminally targeted guide RNA (sgRNA) (gtggtggttaaaccatgtta) was generated using the GeneArt Precision gRNA synthesis kit (Thermo Scientific). Gene editing of individual clones was verified by sequencing with Illumina MiSeq Nano v2, paired end 151 reads, and Sanger sequencing, and clones were additionally validated by immunoblotting and/or mass spectrometry.

**Spinning disk confocal microscopy**

For immunofluorescence staining, cells were fixed with 8% paraformaldehyde in PBS added to an equal volume of media in each well for 4% final paraformaldehyde for 20min at 37ºC, then permeabilized with 0.5% Triton X-100 in PBS for 20-30min at room temperature. Cells were blocked with 3% BSA in PBS with 0.1% Triton X-100 for 1h at room temperature. Cells were incubated with primary antibodies (1:100-1:200 dilution, depending on the primary antibody) in 3% BSA in PBS with 0.1% Triton X-100 for 14-18h at 4°C. After three washes with 0.02% Tween-20 in PBS (TBST), cells were incubated with Alexa Fluor secondary antibodies (1:400) for 1h at room temperature and washed once with TBST, and nuclei were stained with DAPI (1µg/mL final) for 10 min at room temperature. Cells were washed three times with TBST and maintained in PBS at 4ºC until microscopy analysis.

Cells were imaged using a Yokogawa CSU-X1 spinning disk confocal on a Nikon Eclipse Ti-E motorized inverted microscope, which was equipped with a Nikon Plan Apochromat 100×/1.45 NA oil-objective lens and 405- (80 mW), 488- (80 mW), and 640-nm (60 mW) laser lines controlled by a Nikon LUN-F XL solid state laser combiner. Images were acquired with a Hamamatsu ORCA-Fusion BT CMOS camera (6.5 µm2 photodiode, 16-bit) and NIS-Elements image acquisition software. Each channel corresponding to one primary and secondary antibody combination was measured under the same exposure time and laser power across all samples. Z-stacks were collected at 0.5 µm step size.

Fiji/ImageJ software (5) was used to display the z-series as maximum intensity projections and split into channels. The maximum intensity projections were segmented for neuronal soma using CellProfiler Image Analysis Software (Version 4.2.6.) (6). The "Identify Primary Objects" module was used to segment nuclei from the 405-nm/DAPI channel using Otsu threshold (smoothing scale 1.3488, threshold correction factor 0.8, diameter 100-200, size of adaptive window 200, distinguish clumped objects by shape, draw dividing lines by intensity, fill holes after declumping only). Each segmented nucleus was dilated using "Disk", size 50 as structuring element to create a soma mask, which was exported as a binary 8-bit image. Colocalization analysis was performed with BIOP JACoP plugin for Fiji/ImageJ (5) using the masked maximum intensity projections and Yen threshold.

**SDS-PAGE and immunoblotting**

Samples were mixed with 4xLaemmli buffer or 5xLDS buffer, incubated at 80°C for 5 min, and loaded in 4-20% Criterion TGX Stain-free Precast gels for subsequent immunoblotting. After electrophoresis, gels were scanned for total protein using a ChemiDoc MP imager (Bio-Rad) and electro-transferred onto a PVDF or nitrocellulose membrane in Tris/glycine/methanol transfer buffer for 1h at constant current and starting at approximately 45V. Membranes were blocked with 5% non-fat milk or 3% BSA in TBST (0.1% Tween-20), incubated with primary antibody (at 1:500-1:1,000 in 5% non-fat milk or 3% BSA in TBST) overnight at 4°C, washed six times with TBST, and subsequently incubated with HRP-conjugated secondary antibodies (at 1:5,000-1:10,000 in 5% non-fat milk in TBST) for 1h at room temperature. After washing four times in TBST, blot images were acquired using a ChemiDoc MP imager with Western Lightning Plus Chemiluminescence substrate (Revvity, catalog number NEL104001EA). Images were processed with Image Lab software (Bio-Rad, version 6.1.0).

**Endosome and lysosome immunoprecipitation**

For Endo-IPs, hESCs and iNeurons were seeded in 3x15-cm Matrigel-coated dishes per replicate, with iNeurons being seeded on day 4,5, or 6 of the differentiation. For Lyso-IPs, iNeurons were seeded in 2x15-cm Matrigel-coated dishes per replicate. iNeurons were harvested on day 21 of the differentiation, and hESCs were harvested on the same day for consistency. Cells were collected on ice with an initial on-plate wash using 2mL cold PBS, then scraped and transferred to tubes using wide-bore transfer pipettes and 1mL PBS supplemented with protease inhibitors (Roche). Residual cells were recovered from the plates with an additional 1mL of PBS with protease inhibitors. Cells were spun down at 500xg for 4min at 4ºC, supernatant was removed, and cells were gently resuspended in 950μL of KPBS (100mM potassium phosphate, 25mM KCl, pH 7.2) with protease inhibitors (KPBS+i). Cells were lysed on ice with 30 strokes of a 2mL Dounce homogenizer (DWK Life Sciences) and B type pestle. Lysates were pelleted at 1,000xg for 5min at 4ºC, supernatants were transferred to new tubes, and samples were spun again at 1,000xg for 5min at 4ºC. The final post-nuclear supernatants (PNS) were transferred to new tubes on ice. Protein quantification of each PNS was measured by Bradford. Protein concentration of each PNS was normalized based on the Bradford results, and 10µL of each sample was combined with 30µL RIPA buffer and 10µL 5xLDS buffer and DTT and set aside for immunoblotting. For Endo-IPs, anti-FLAG M2 magnetic beads (Sigma-Aldrich) were washed three times with KPBS+i, and 70µL of bead slurry was prepared per lysate. For Lyso-IPs, anti-HA magnetic beads (Pierce) were washed in the same manner, and 65µL of bead slurry was prepared per lysate. Normalized PNS samples were incubated with beads at 4ºC with gentle rotation for 30min for Lyso-IP and 50min for Endo-IP. After incubation, 10μL of each flow-through was combined with 30μL RIPA and 10μL 5xLDS and DTT and set aside for immunoblotting. Anti-FLAG beads were washed three times with KPBS+i, and anti-HA beads were washed twice with high salt KPBS+i (supplemented with 150mM NaCl) and once with KPBS+i. Samples were eluted with 120µL 0.5% NP-40 in KPBS+i at 4ºC with gentle rotation for 30min. For analysis by immunoblotting, 20µL of each eluate was combined with 6µL 5xLDS and DTT, and 100µL of each eluate was further processed for mass spectrometry.

**Proteomics**

**S-trap sample preparation and TMT labeling**

Eluate samples from Endo-IP and Lyso-IP were mixed with 100µL of water, 200µL lysis buffer (10% SDS, 100mM Triethylammonium bicarbonate (TEAB) pH 8.5) and processed using S-Trap micro-spin columns following the manufacturer's protocol (Protifi, C02-micro-80) (7, 8). Proteins were reduced with 5mM TCEP and incubated at 55ºC for 30min with shaking at 800rpm. Cysteines were alkylated with freshly prepared chloroacetamide at 40mM final concentration at 25ºC for 30min with shaking at 800rpm and protected from light. Proteins were digested with 0.5µg Lys-C at 37°C overnight then with 0.5µg of trypsin for 6h with shaking at 800rpm. Peptides were collected and dried in a SpeedVac.

Peptides were resuspended in 35µL 100mM TEAB pH 8.5 and 11µL ACN and labelled with 10µL of TMTpro reagent (20 mg/mL stock) for 1h at room temperature. The reaction was quenched with 10µL 5% hydroxylamine for 15min. Equal peptide amounts of each sample were combined and fractionated using High-pH Reversed-Phase Peptide Fractionation Kit (Pierce). Six fractions were collected and desalted with C18 StageTips. Fractions were reconstituted in 5% ACN, 5% formic acid and analyzed by LC-MS/MS.

**Liquid Chromatography-Mass spectrometry (LC-MS) analysis**

Lyso-IP and Endo-IP samples (**Dataset S1**) were analyzed in an Orbitrap Eclipse Tribid mass spectrometer coupled to a Vanquish Neo UHPLC system. Endo-IP samples from hESC and iNeurons (dataset 2) were analyzed in an Orbitrap Fusion Lumos mass spectrometer coupled to an EASY-nLC 1200 system. Peptides were separated on a 100µm microcapillary column packed with 20cm of Accucore C18 resin (2.6 µm, 150 Å) with a 90-min linear gradient from 5% to 30% ACN in 0.125% formic acid. FAIMS Pro was set to -30, -50 and -70 compensation voltage (CV). MS1 spectrum was acquired on the Orbitrap (60,000 resolution, 350-1,350 m/z, standard automatic gain control (AGC) target, auto maximum injection time). MS2 was acquired on the Orbitrap (30,000 resolution, HCD at 36% NCE, 0.6 m/z isolation window, TurboTMT set to All TMT Reagents, 200% normalized AGC, 120ms maximum injection time).

TMT-MS data was analyzed with MSconverter (9) and Comet (10). Peptide mass tolerance and fragment ion tolerance were set to 50ppm and 0.02 Da, respectively (9-11). Peptide modifications included: TMTpro labels as fixed modification (+304.207 Da on lysines and peptide N-terminus), carboxyamidomethylation as fixed modification (+57.021 Da on cysteines), and oxidation as variable modification on methionines. Peptide-spectrum matches (PSMs) were filtered by linear discriminant analysis (11) to 2% false discovery rate (FDR)(12). TMT-reporter ions were quantified and corrected for isotopic impurity. PSMs were filtered for at least 200 total signal-to-noise ratio and 50% precursor isolation purity.

**Candidate cargo analysis and Structural predictions**

Endosomal proteins were annotated according to previous experimental data and literature, such as (13, 14). Proteins were considered candidate endocytic cargo if they were: 1) significantly enriched in our Endo-IP from iNeurons, 2) contained at least one transmembrane segment, and 3) were annotated as localized to the plasma membrane (based on Uniprot). SNX17 motifs were searched only in cytoplasmic regions for candidate endocytic cargo: Φ-x-N-x-x-[FY] and N-P-x-x-[FY]. The SNX27 motif searched was: [S/T/D/E]-[S/T]-x-Φ-C-term, where Φ is A/V/I/L/M/F/Y/W. The results of searches are provided in **Dataset S3**. The binding of candidate cargo with SNX17 and SNX27 were modelled using 15 residue-long peptides containing the SNX motifs in AlphaFold 3, as described (15), and visualized using Pymol (Version 2.5.4).

**Statistics and reproducibility**

For MS experiments, 3 to 4 biological replicates per condition were performed and analyzed using MSstats (16, 17). Statistical analysis was obtained with peptide level normalization and imputation enabled, and MSstats protein summarization. Proteins were considered significantly enriched in Endo-IP with $<0.01$ q-value and $>2$ fold-change. Synaptic Gene Ontology enrichment analysis was performed using SynGO (18) (https://www.syngoportal.org/#) with all proteins identified in each experiment as background. For immunofluorescence experiments, at least 3 technical replicates (where each replicate represents an individual well in a 24W glass-bottomed plate) per cell line per antibody combination were imaged and analyzed, with multiple images taken per replicate well. Two-sided unpaired T-tests were performed, and p-values were reported.

### Supplemental Figures

**Fig. S1**

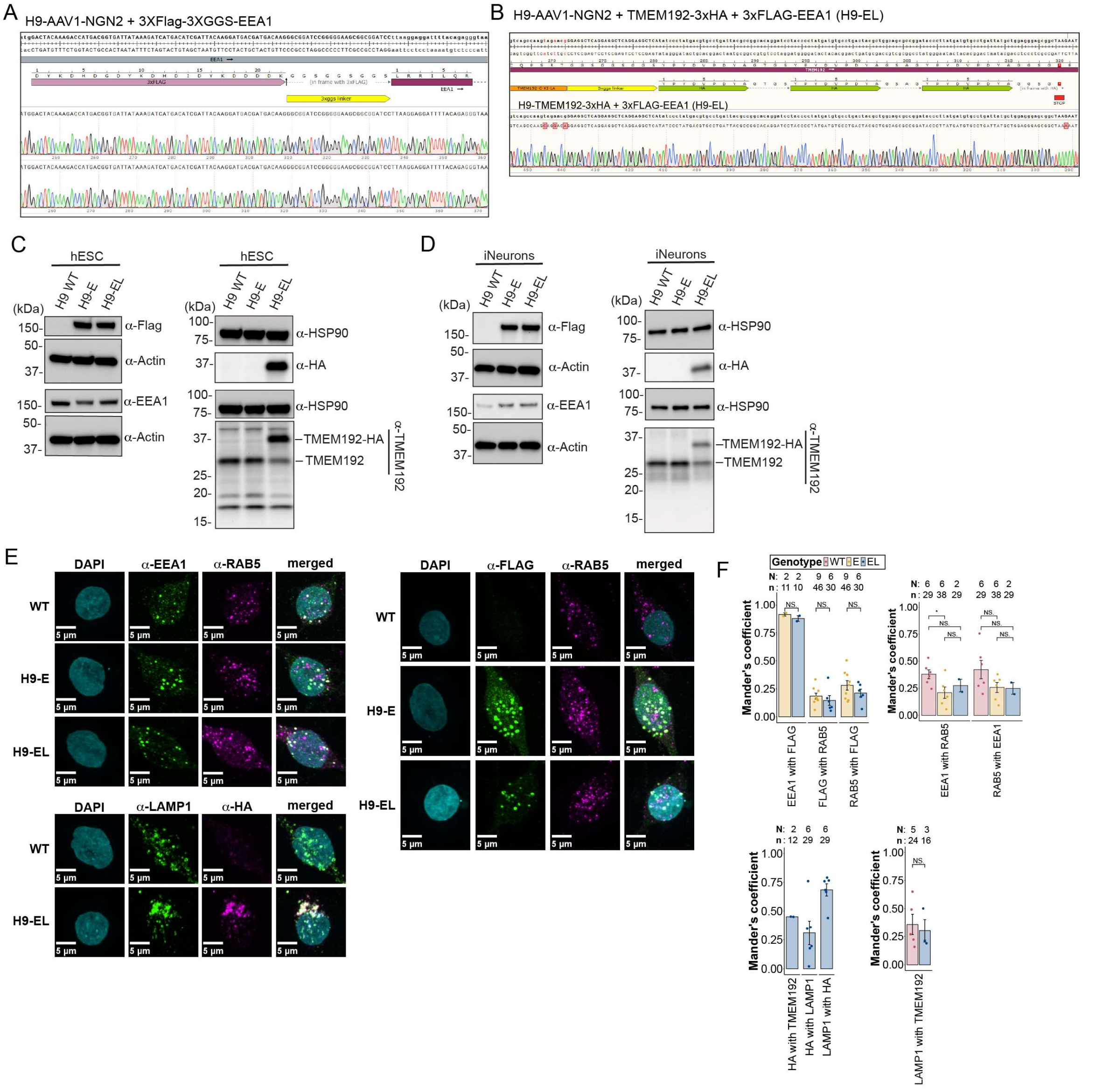


**Fig. S1. Toolkit and validation of Endo-IP and Lyso-IP for organelle proteomics in hESC-derived iNeurons. (A)** Sanger sequencing analysis of H9-E (top) and H9-EL (bottom) clones after gene editing with 3xFLAG at the N-terminus of EEA1. **(B)** Sanger sequencing analysis of the H9-EL clone showing gene editing with 3xHA at the C-terminus of TMEM192. Downstream NeoR not depicted. **(C)** Western blots of hESC whole cell lysates from H9 WT untagged, H9-E, and H9-EL cells showing FLAG tagging of EEA1 in both E and EL cells and heterozygous HA tagging of TMEM192 in EL cells. **(D)** Western blots of day 21 iNeuron whole cell lysates from H9 WT untagged, H9-E, and H9-EL. **(E)** Confocal microscopy max intensity projections of day 21-23 iNeuron somas showing α-EEA1, α-RAB5, α-LAMP1, and α-HA staining in H9 WT untagged, H9-E, and H9-EL cells. Images are representative examples. **(F)** Quantification of mean Mander's coefficients from (E). Each dot represents the mean Mander's coefficient per well imaged (with three or more images per well), and error bars represent standard error of the mean. *P ≤ 0.05, two-sided unpaired T-test. N=the number of replicates per combination, where each well imaged is a replicate; n=the number of images per replicate.

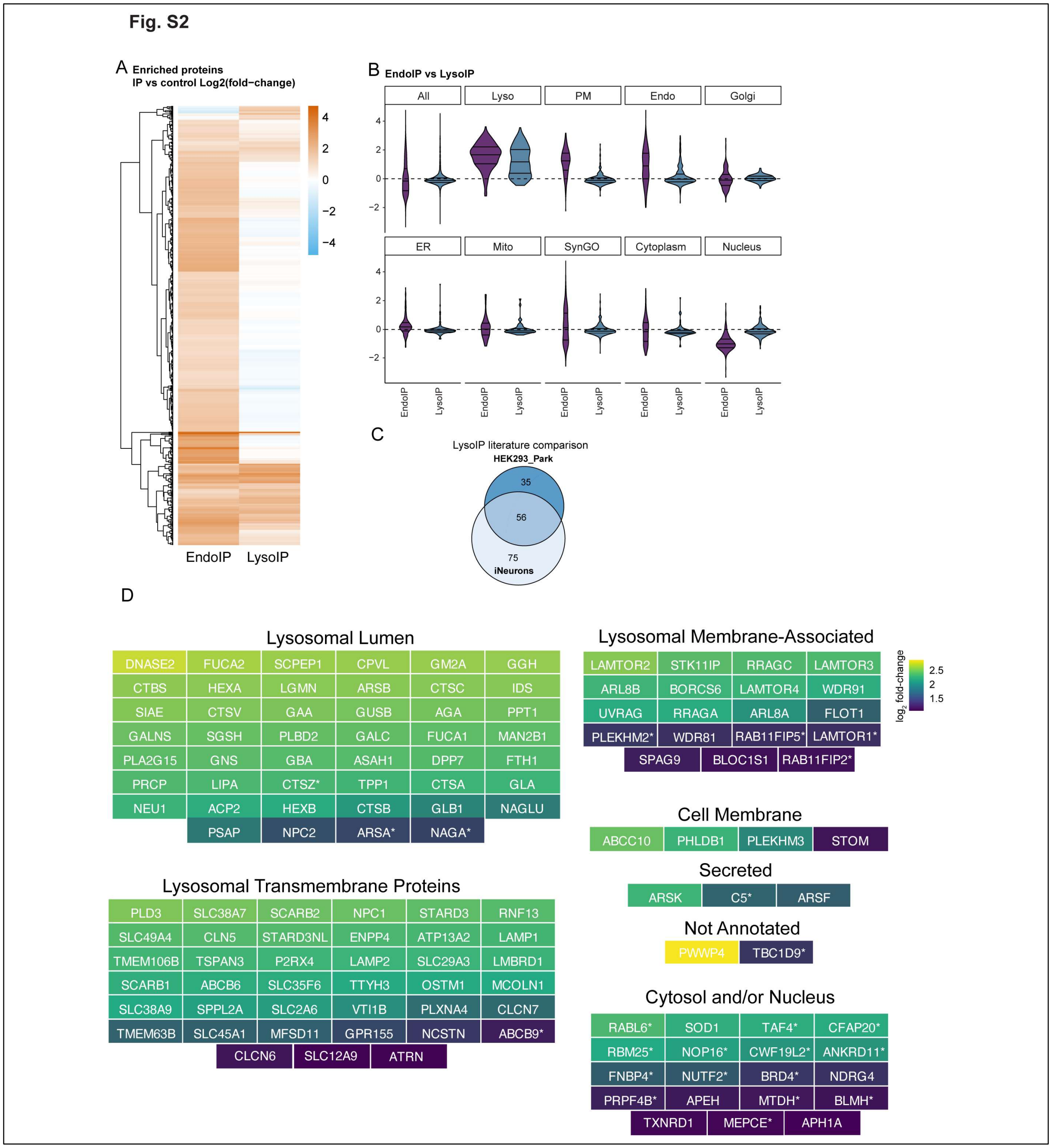


**Fig. S2. Proteomic landscape of EEA-positive endosomes in hESCs and iNeurons. (A)** Heatmap showing the abundance fold-changes ($\log_2$) of all proteins detected in both the Endo-IP and Lyso-IP from day 21 iNeurons. **(B)** Violin plots depicting the enrichment of proteins of different subcellular localizations (see **Materials and Methods**) and SynGo annotation in Endo-IP (purple) and Lyso-IP (blue) normalized to untagged controls. **(C)** Comparison of all proteins significantly enriched ($Log_2FC \geq 1.0$, q-value $\leq 0.01$) in Lyso-IPs from iNeurons (this study) versus proteins significantly enriched ($Log_2FC \geq 1.0$, p-value $\leq 0.02$) in Lyso-IPs from 293 cells (19). **(D)** Sublysosomal and subcellular localization of all proteins significantly enriched (Log$_2$FC ≥ 1.0, q-value ≤0.01) in Lyso-IPs from iNeurons. Many proteins (111 out of 132) were also enriched in Endo-IPs from iNeurons, and proteins exclusively enriched by Lyso-IP are indicated by asterisks.

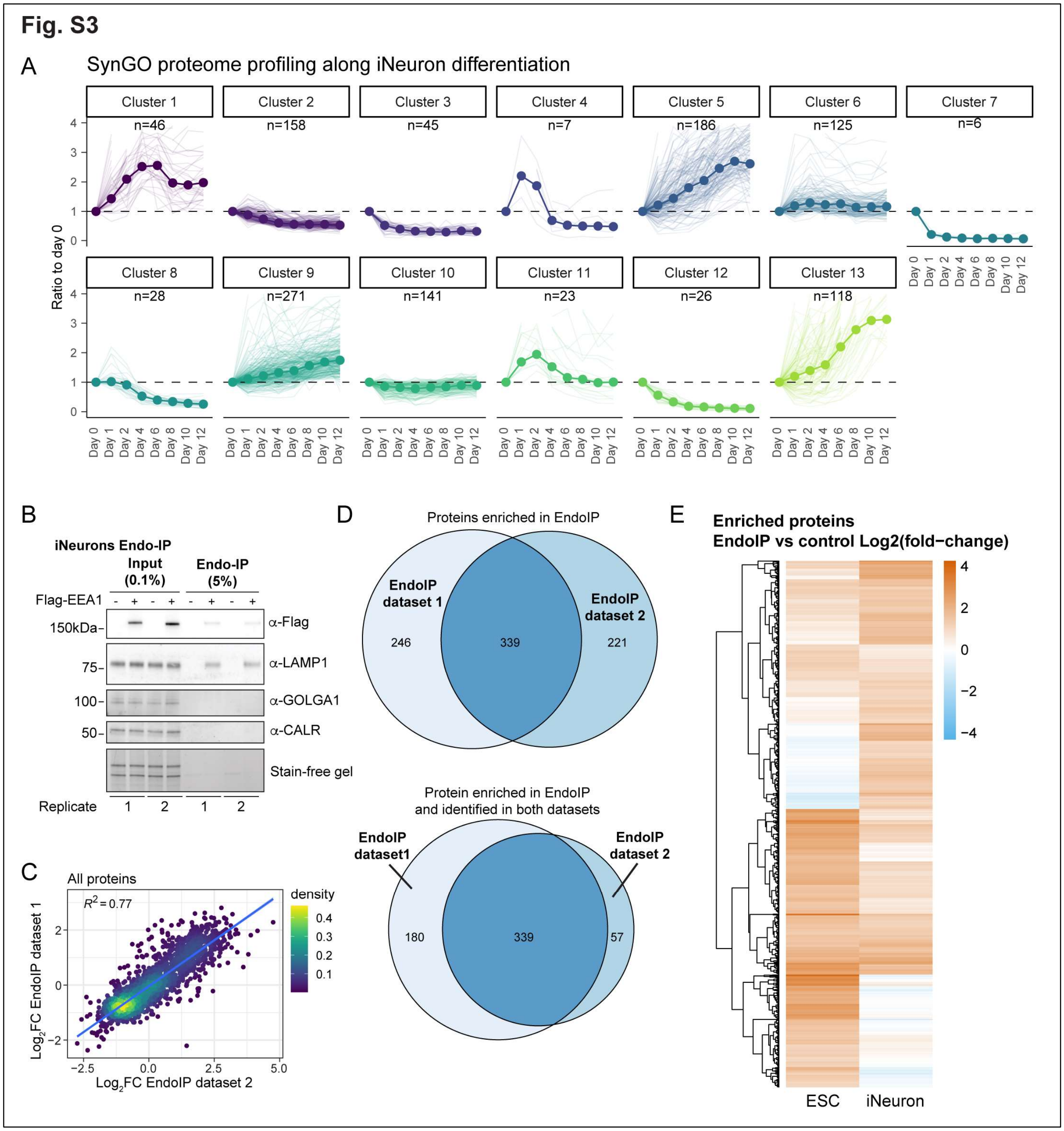


**Fig. S3. Repertoire of receptor and synaptic protein cargo in iNeurons. (A)** Cluster analysis of all SynGo-annotated proteins detected over differentiation of iNeurons (20) where several of the largest clusters (e.g. clusters 5, 9, and 13) show increased expression over time. **(B)** Western blot of input and eluate samples from iNeuron experiment in Fig. 2C (two example replicates are shown) showing α–FLAG, α–LAMP1, α–GOLGA1 (Golgi), and α–CALR (Endoplasmic Reticulum). **(C)** Correlation between Endo-IPs from **Fig. 1B** (**Dataset S1**) and **Fig. 2C** (**Dataset S2**), depicting only proteins that were detected in both experiments. **(D) Top:** Comparison of proteins significantly enriched (Log$_2$FC ≥ 1.0, q-value ≤0.01) in Endo-IPs from Fig. 1B (**Dataset S1**) and **Fig. 2C** (**Dataset**

S2). **Bottom:** Comparison of proteins significantly enriched (Log$_2$FC ≥ 1.0, q-value ≤0.01) and detected in both datasets. **(E)** Heatmap showing the abundance fold-changes (Log$_2$) of all proteins detected in Endo-IPs from both hESCs and day 21 iNeurons.

**Fig. S4**

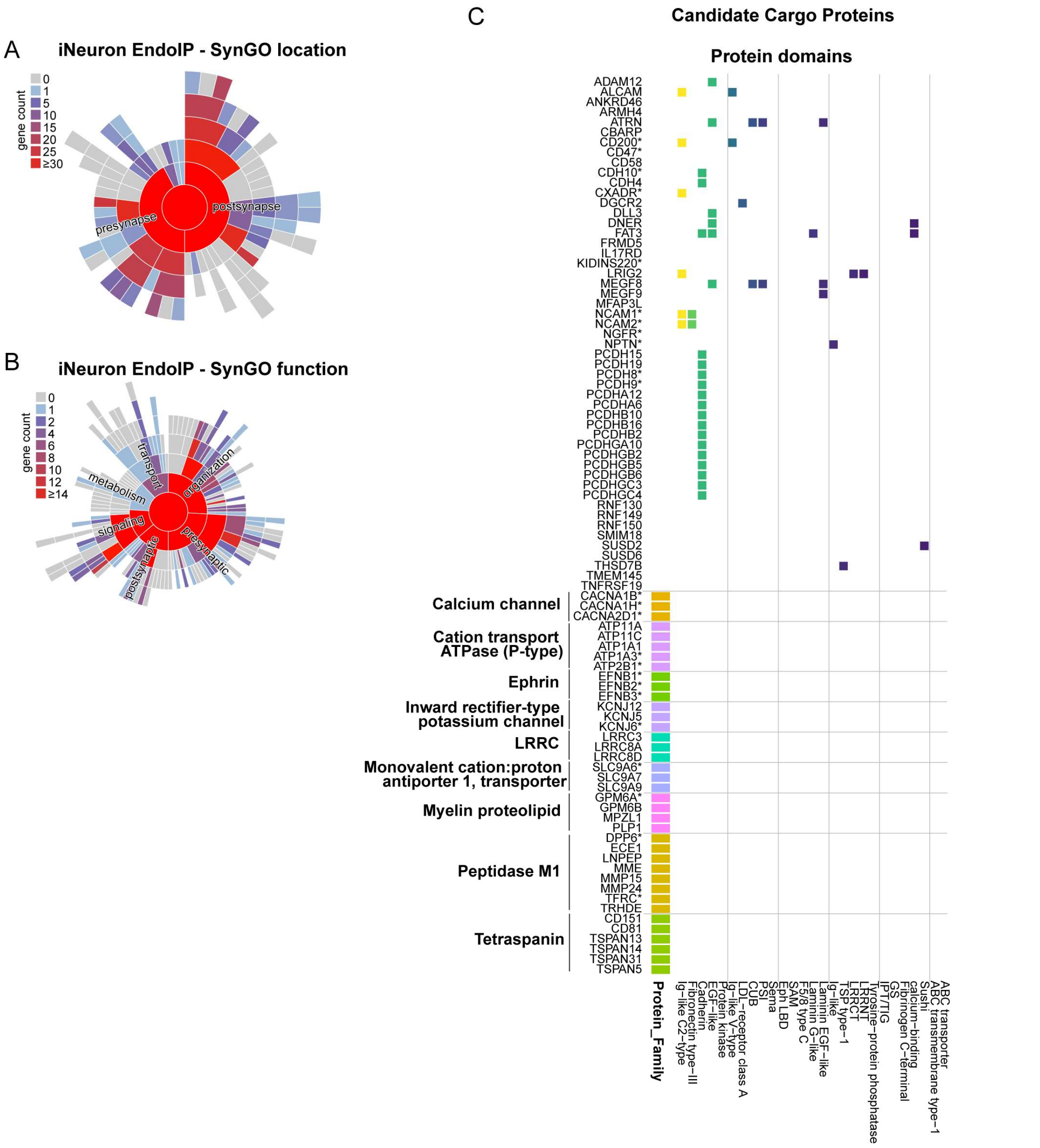


**Fig. S4. Identification of candidate neuronal SNX27 and SNX17 cargo. (A)** SynGo location analysis of proteins significantly enriched (Log$_2$FC ≥ 1.0, q-value ≤0.01) in Endo-IPs in day 21 iNeurons. **(B)** SynGO function analysis of proteins significantly enriched (Log$_2$FC ≥ 1.0, q-value ≤0.01) in Endo-IPs from day 21 iNeurons. **(C)** As in **Fig. 3E**, analysis of candidate cargo proteins significantly enriched in Endo-IPs from day 21 iNeurons which harbor one or more transmembrane segment and are annotated as localized to the cell membrane, and which have either an annotated protein family or domain, but not both.

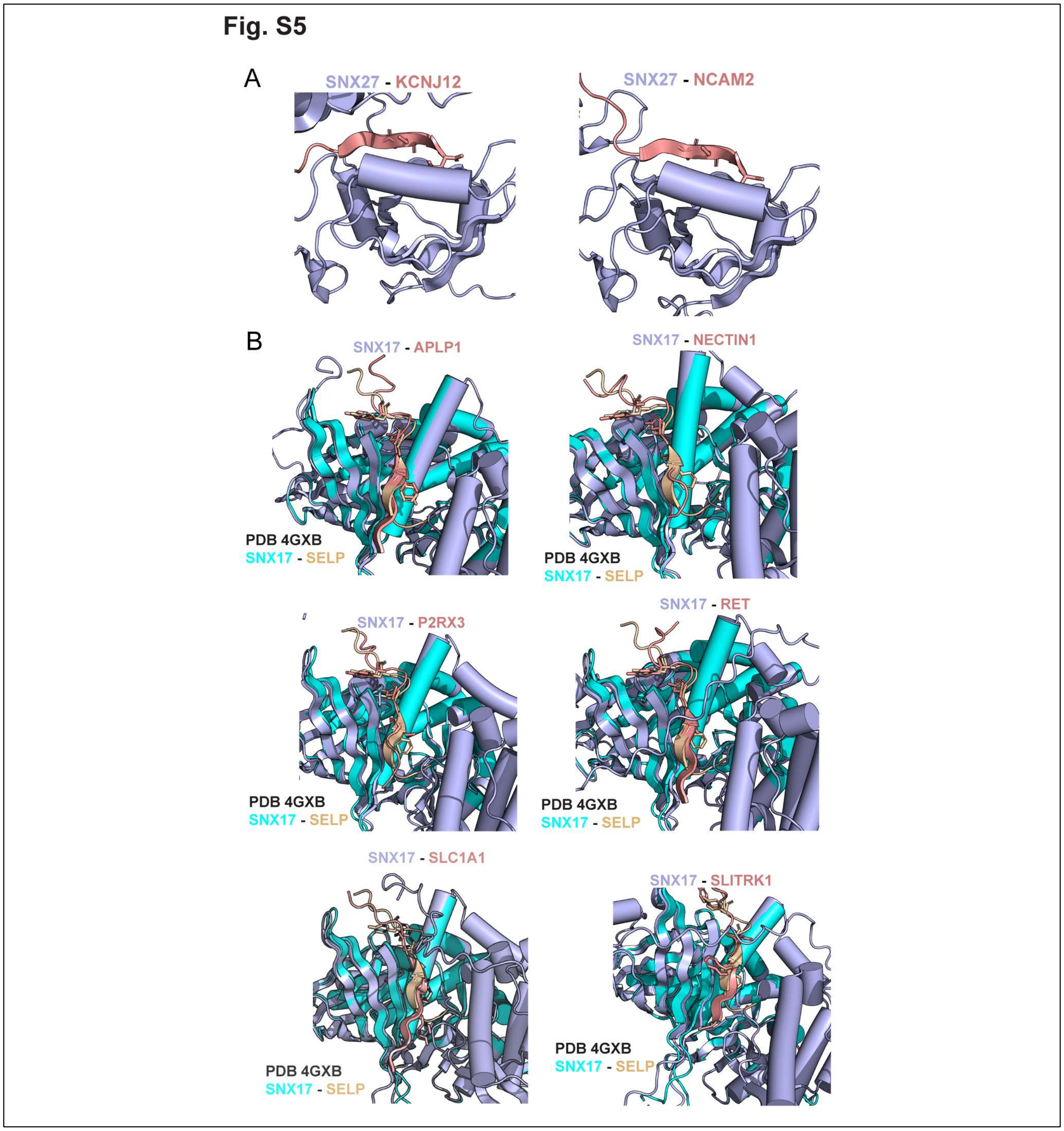


**Fig. S5. Matching sorting motifs with candidate cargo. (A)** AF3 predictions of SNX27 and 15 amino acid peptides containing a putative SNX27 binding motif in KCNJ12 and NCAM2. **(B)** Overlay of SNX17-SELP structure (PDB: 4GXB, in cyan and brown) with AF3 predictions for SNX17 (in light blue) bound to 15 amino acid peptides containing putative SNX17 binding motifs from APLP1, NECTIN1, P2RX3, RET, SLC1A1, and SLITRK1 (in salmon). The SELP residues FxNxxY shown to mediate binding to SNX17 are depicted as sticks, as are the residues predicted to mediate binding in the six candidate cargo proteins.

**Dataset S1 (separate file).** Proteomic analysis of Endo-IP and Lyso-IP from day 21 iNeurons.
**Dataset S2 (separate file).** Proteomic analysis of Endo-IPs from hESCs and day 21 iNeurons.
**Dataset S3 (separate file).** SNX27 and SNX17 sorting motif predictions for candidate endocytic cargo from iNeurons.
